# Human endothelial cells display a rapid and fluid flow dependent tensional stress increase in response to tumor necrosis factor-*α*

**DOI:** 10.1101/2022.01.12.476017

**Authors:** Matthias Brandt, Volker Gerke, Timo Betz

## Abstract

As endothelial cells form the inner layer of blood vessels they display the first barrier to interstitial tissues, which results in a crucial role for inflammation. On the global, systemic level an important element of the complex process controlling the inflammatory response is the release of the cytokine tumor necrosis factor-*α* (TNF-*α*). While other pro-inflammatory agents like thrombin or histamine are known to induce acute but transient changes in endothelial cells which have been well studied biologically as well as mechanically, TNF-*α* is primarily known for its sustained effects on permeability and leukocyte recruitment. These functions are associated with transcriptional changes that take place on the timescale of hours and days. Here we show that already 4 minutes after the addition of TNF-*α* onto monolayers of human umbilical vein endothelial cells, a striking rise in mechanical substrate traction force and internal monolayer tension can be recorded. As expected, the traction forces act primarily at the boundary of the monolayer. While the traction forces increase monotonically during the initial cellular response, we find that the internal monolayer tension displays a rapid peak that can be abolished when applying a shear flow to the cells. The increased internal monolayer tension may provide a mechanical signal for the cells to prepare for the recruitment of leukocytes, additionally to the well studied biochemical response.

## Introduction

Endothelial cells (ECs) constitute the interior surface of blood vessels and form the key barrier to the surrounding tissue, ensuring among other things that blood cells do not undesirably enter. In this function, ECs are known to regulate the exchange of molecules and fluid, but also of immune cells and even metastatic cancer cells in pathological situations. This gatekeeper function plays hence an important role in physiological maintenance not only of the vascular system, but also of the overall tissue of any organism. Since ECs are constantly challenged by mechanical stress due to blood pressure, surrounding tissue stiffness and vessel stretch, the importance of mechanical stress in vascular diseases has gained increasing attention over the last few decades and is well established by now [1, 2]. While many inflammatory processes are known to increase mechanical tension and barrier permeability of these key cellular monolayers [2, 3], it is also proposed that the mechanical force load on the ECs can in itself alter the immune response and affect proper guidance and migration of immune cells into injured tissues [4, 5]. In stark contrast to this well described importance of mechanical forces and endothelial mechanics for the proper immune response, we have only sparse knowledge about the early mechanical response of endothelial cells to inflammatory signals such as tumor necrosis factor-*α* (TNF-*α*). Although a rapid chemical reaction of ECs to TNF-*α* is known, the link to a mechanical reaction has only been established in the long time limit. TNF-*α* is one of the central pro-inflammatory signaling molecules and is mostly supplied by macrophages and monocytes upon their activation via pathogen interaction. A prominent function of this cytokine is to increase endothelial permeability and to sustain enhanced leukocyte recruitment [6–8]. While these effects are mostly associated with transcriptional changes, thus taking place on a slow timescale of several hours, other pro-inflammatory agents like thrombin or histamine are known to induce acute transient changes in ECs that take place within minutes [9, 10]. Biophysical studies measuring either substrate traction forces exerted by ECs on elastic hydrogels or monolayer tension using isometric force transducers could confirm a quick and strong enhancement of EC monolayer contractility associated with increased cell-cell forces in response to Thrombin [11, 12]. In case of TNF-*α* on the other hand, biophysical studies focused mostly on long term changes in the range of 4-24 hours after exposure involving reduced cortical stiffness, cell elongation and enhanced cell-substrate forces [13–16]. Partially, these studies were carried out on single cells only and not on interconnected monolayers of ECs. However, rapid morphological changes similar to the response to thrombin, though weaker, have also been reported for TNF-*α*. TNF-*α* was found to induce actin cytoskeletal reorganizations within the first 5-10 minutes of stimulation, i.e. enrichment of fine actin cables and formation of lamellipodia and filopodia [17]. After 15-30 minutes these changes are followed by the increased formation of stress fibers. At the same time ECs begin to retract and VE-cadherin starts to dissociate from intercellular junctions. The formation of stress fibers in response to TNF-*α* has been linked to increased levels of RhoA activity promoting myosin light chain (MLC) phosphorylation via downstream activation of Rho-associated kinase (ROCK) [18, 19]. MLC phosphorylation in turn is well known to increase actomyosin contractility [20]. However, the mechanical response in form of force generation onto the substrate and also on the cell neighbours remains to be explored. Recently, not only the mechanical interaction with the substrate has been shown to be important for the mechanical equilibrium of cell layers, but also the force transmission to the neighbouring cells via cell-cell adhesion has become of increasing interest to better understand the biomechanics of cell layers in general [21–23] and endothelial cells in particular [11, 24–26]. To describe the direct mechanical interaction between ECs, their substrates and their neighbours, we use patterned monolayers combined with traction force microscopy to gain access to both the substrate traction and the cell-cell forces exerted by endothelial cells.

## Results and Discussion

### Circular HUVEC monolayers apply substrate stress primarily at their boundary

To determine the mechanical response of a human umbilical vein endothelial cell (HUVEC) monolayer upon the exposure of TNF-*α*, traction force microscopy was performed on cells seeded overnight on a polyacrylamide (PAA) gel of 15 kPa stiffness and grown in circular patches. Micropatterning of collagen coating of the gel surface led to the formation of circular shaped monolayers with a diameter of 300 *μ*m (Fig. 1A). The size of the pattern was chosen to allow for the formation of bigger cell clusters that resemble a monolayer, while still keeping the pattern small enough to fit into the microscope’s field of view using a 40X water immersion objective (for initial experiments on stained cells shown in Fig. 1 images were recorded with a 60X objective with higher numerical aperture instead and stitched together for better quality). The pattern allows to differentiate between the mechanical forces acting on a substrate in the region of intact monolayer, but also at the edge regions of the pattern, where the monolayer is disrupted and an open edge exists. Consistent with reports on epithelial cells [27, 28], we find the traction forces to be focused at the monolayer edge (Fig. 1B). To quantify this, we compared average forces exerted by cells at the edge and in the inner region of the pattern. The difference between edge and inner cells was defined by the manually measured average cell diameter of edge cells of about 19 *μ*m. Using this radial distance from the pattern edge, we find that with an average of ≈ 198 Pa edge cells exerted 65 % higher traction stresses than inner cells which only exhibited an average of ≈120Pa (Fig. 1C). Noteworthy, the background level already lies at about 100 Pa making this discrepancy even more striking. The reason for such a behaviour in general can be explained by two hypotheses. First, it is possible that in fact only the cells at the periphery are contractile, and hence only these are applying a reasonable force on the substrate. Alternatively, in a monolayer forces can be passed from cell to cell by intercellular junctions, in particular adherence junctions. In this scenario the cells at the edge do simply lack neighboring cells to further connect to at the patterns periphery, and are therefore required to transmit the forces they received from their inner neighbours onto the substrate to achieve force balance. This second explanation has been demonstrated to be correct in other monolayer systems previously [27] and was hence tested for the here presented system.

**Fig 1.**
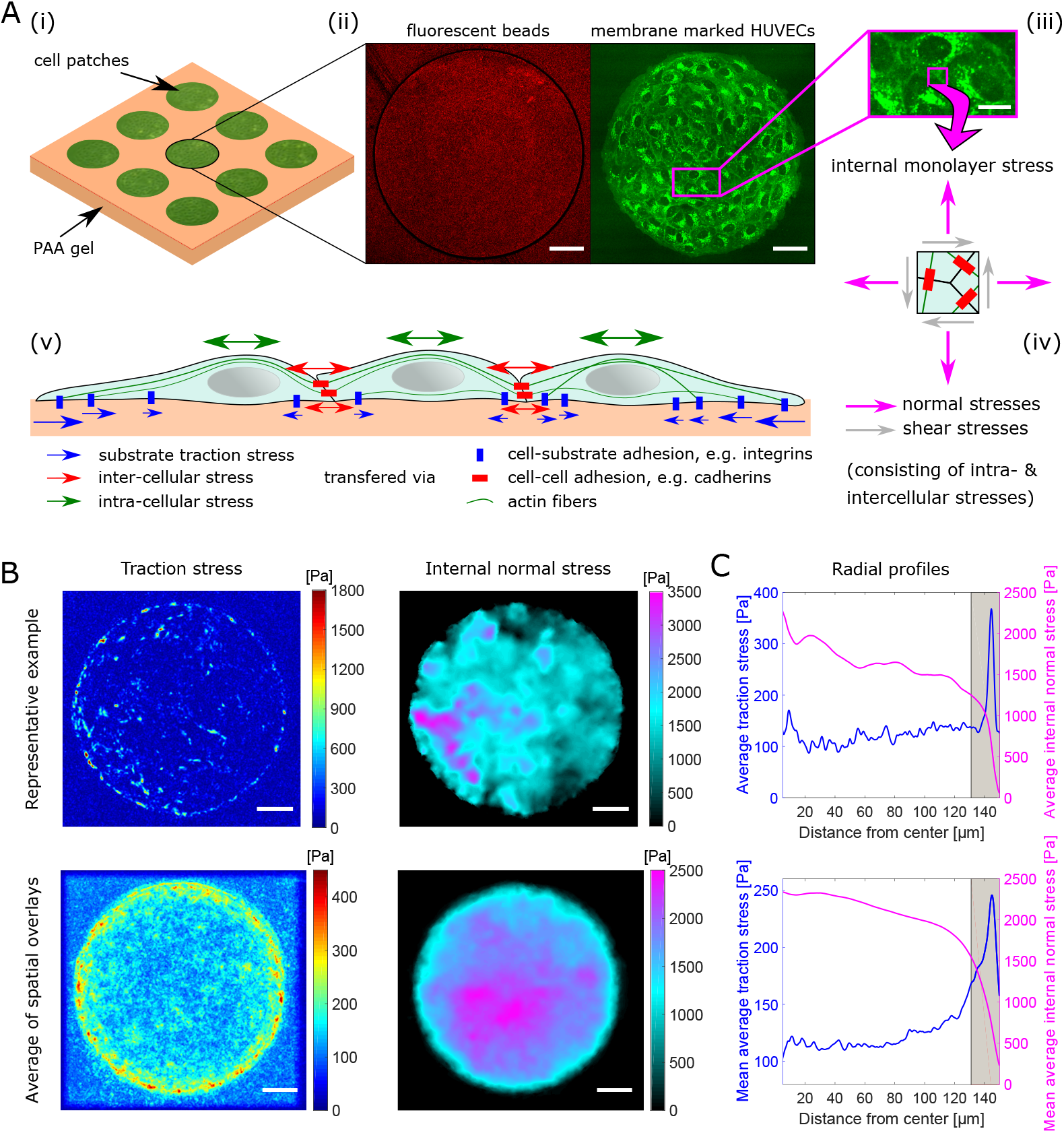
Experimental system and mechanical stress distributions. **A**: Circular islands of EC monolayers reside on micropatterned PAA gels (**i**) that incorporate fluorescent beads (**ii** - red) close to the surface for traction force microscopy. (**iii**) Zoom in on plasma membrane stained (CellMask Deep Red) HUVECs in a monolayer patch (**ii** - green). (**iv**) Normal and shear stresses act at any position within a monolayer of cells. (**v**) Schematic illustration of mechanical stress exerted and transmitted by interconnected cells. Scale bars, 50*μ*m (15*μ*m for zoom in). **B**: Color coded maps of derived traction stress and internal normal stress for a representative example monolayer patch (top) and for the spatially overlayed and averaged total of 22 different patches from 3 independent experiments (bottom). Scale bars, 50 *μ*m. **C**: Corresponding radial profiles of average traction and internal normal stress based on the data used in B (top for single example, bottom for average of multiple patches). The gray marked area indicates the approximate region of edge cells. The inner 5 *μ*m have been neglected here due to poor number of pixels statistics. Traction stresses are applied dominantly by cells at the edge of the monolayers while internal normal stresses accumulate towards the center.

### Transcellular stress transmission in the monolayer region dominates traction forces

To assess stresses passed from cell to cell we employed monolayer stress analysis similar as introduced previously [21, 24]. In short internal monolayer stresses are inferred from measured traction forces by imposing force balance under the assumption of homogeneous material properties throughout the whole cell monolayer. As expected and consistent with previous reports on other monolayer systems we find high internal monolayer stress transmission throughout the whole circular pattern, with a high stress accumulation at the central region of the pattern (Fig. 1B). The latter might be a geometrical effect that arises from the shape of the pattern and is not necessarily reflecting the real stress distribution throughout a HUVEC monolayer in general. The internal stress maps show here the average of only the normal component which in general represents either tensional (positive values) or compressive (negative values) stresses as opposed to shear stresses that run parallel to a surface (see Fig. 1A-(iv)). However, comparing again average values for edge versus inner cells the picture is inverted for the tensional monolayer stress with respect to the traction forces. Here we find two times larger average stresses of ≈2.1 kPa for inner cells as compared to ≈1.0kPa for cells residing at the edge of the pattern. The observation of low substrate traction forces but high internal monolayer stresses in the central region and vice versa for the outer region of the monolayer is best explained by a high transcellular stress transmission through the monolayer. That the difference between outer and inner cells is already striking after one cell diameter of 19 *μ*m and plateaus at about 50 *μ*m away from the edge of the pattern further suggests that endothelial cells actually prefer to transmit stress among themselves instead of applying forces on the substrate. However, the precise force transmission that is expected to be passed from cell to cell via stress fibers remains undetected in this approach. An interesting speculation is that the generation of defects or holes in a monolayer, for example during high stress within a blood vessel or because of toxins, leads to a shift of force transmission from cell-cell contacts to substrate contacts. Such changes in force transmission, which are a purely mechanical effect and independent of molecular signaling, would effectively allow a cell to recognise monolayer rupture by simple engagement of substrate forces.

### TNF-alpha increases traction forces and transcellular stress

Having established a method to determine both the traction forces and the internal monolayer stresses of an endothelial cell layer, we went on to investigate potential changes of the mechanical properties of HUVEC monolayers in response to TNF-*α* signaling. It is well established that long-term application of TNF-*α* leads to prominent stress fiber formation and increased permeability in EC monolayers. In single cells also a long-term increase of contractility mediated by substrate forces has been found [14]. However, to which extent the overall stress distribution inside a monolayer is affected by TNF-*α* has not been studied yet. As presented in Fig. 2B, after applying TNF-*α* for 1 hour at 20 ng/ml concentration we find a significant increase of both, average substrate traction forces as well as average internal normal stress throughout the monolayer. Both, traction forces and internal stresses increase similarly in magnitude, and no qualitative change in the force distribution is observed. Similar to control conditions, traction forces are displayed predominantly at the edge while normal stresses within the layer built up towards its center Fig. 2A. Together this hints to a joint effort contraction of the whole cell monolayer patch. This is further supported visually by comparing fluorescent images of membrane stained cells in both conditions (S1 Video).

**Fig 2.**
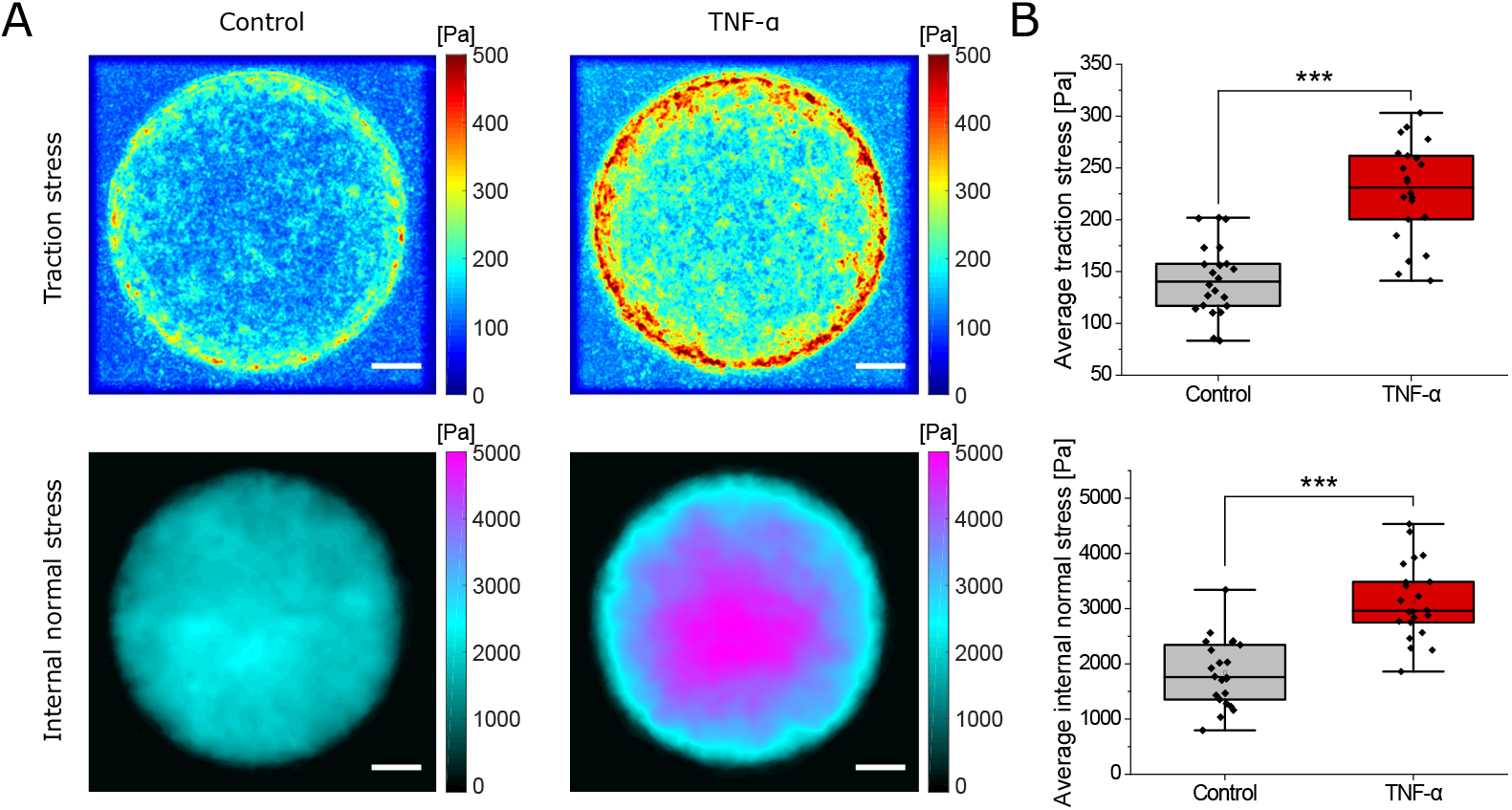
Increase of both traction and internal normal stress upon 20ng/ml TNF-*α* exposure of HUVECs for 1 h. **A**: Spatially overlayed and averaged color coded maps of derived substrate traction and internal normal stress of multiple monolayer patches prior to (Control) and upon application of TNF-*α*. Scale bars, 50 *μ*m. **B**: Average traction stress (top) as well as internal normal stress (bottom) increased significantly after 1 hour of TNF-*α* treatment. Data for A and B is based on *n* = 22 monolayer islands from *N* = 3 independent experiments; ***, *p* < 0.001.

### Rapid increase of substrate traction stress upon TNF-*α* stimulation

To better understand the time scale of the mechanical response we recorded the dynamics of contractility by taking full 3D stacks of cells and substrate deformation for multiple patches with 4 minutes time resolution. To avoid photodamage on the cells due to the prolonged exposure of fluorescent membrane markers, cells were observed using bright-field illumination only. In these experiments, we initially recorded the cells and the deformation for 20 minutes to obtain a force baseline. Subsequently, we added 20ng/ml TNF-*α*. As shown in Fig. 3A,B we find an immediate response as average stresses already started to rise within the first acquired time interval (*p* = 0.0021), with a typical peak at about 20 minutes after applying the chemokine.

**Fig 3.**
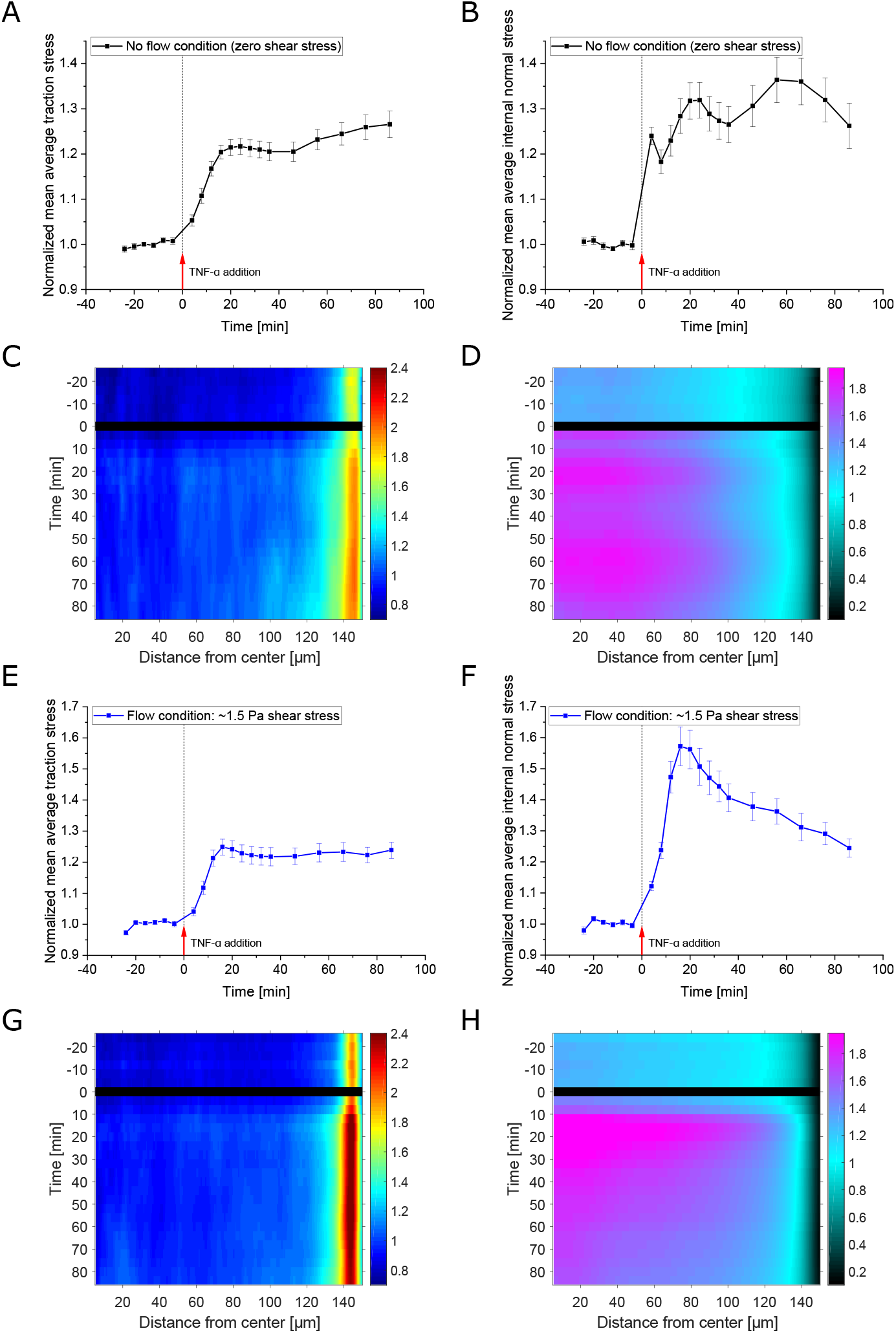
Time evolution of the mechanical response of HUVECs upon TNF-*α* exposure in no flow (zero shear stress) and flow (1.5 Pa shear stress) conditions. **A,B**: Time course of the mean of spatially averaged traction stress (A) and internal normal stress (B) of multiple monolayer islands recorded at baseline and upon 20ng/ml TNF-*α* treatment under no flow (zero shear stress) condition. Data is shown as mean±SEM and has been normalized to the mean of the first 6 timepoints recorded prior to drug administration (*n* = 29, *N* = 3). Average stresses rise immediately upon TNF-*α* treatment (*p* = 0.0021 at 4 min vs −4 min) reaching a peak after about 20 min. While substrate traction stress is increasing monotonically, for the internal normal stress a quick steep rise (*p* < 0.001 at 4 min vs −4 min) with a subsequent intermediate drop (*p* = 0.027 from 4 to 8 min) can be observed. **C,D**: Kymographs showing the development of radial profiles of substrate traction (C) and internal normal stress (D) over time. Kymographs are based on the same measurements as A, B. Colors encode the normalized stress intensity, normalized as in A, B respectively. As application of TNF-*α* required some time and caused disturbance, no image data was analyzed for the respective timepoint 0 (black line). The overall increase in stress intensity fits the time course of the average values shown in A and B. While the rapid increase in traction stress is largely generated at the edge of the monolayer, a delayed increase of traction stress in the inner region can still be observed as well. Interestingly, while there is hardly any increase in the inner region for the traction stress during the first time interval acquired after TNF-*α* application, the internal normal stress of the inner region is highly enhanced already at this point in time (D). **E-H**: Same as in A-D, respectively, but for measurements under the condition of flow induced shear stress (*n* = 35, *N* = 3). The time course of the average traction stress in response to TNF-*α* under flow condition closely follows the one observed in case of no flow. The initial steep rise with subsequent intermediate drop in internal normal stress, however, is not reproduced in the presence of flow and the peak level reached is significantly higher than in situation without applied flow (*p* = 0.0022 at 20min peak). Further, the discrepancy between edge and inner region of the monolayer for the exerted traction stress is higher in the presence of flow (see also S2 Fig.).

### Signs of enhanced transcellular stress transmission and rapid redistribution of stress

As we find that both traction forces as well as internal monolayer stresses increased by the same relative amount after one hour in membrane stained ECs, we expected that the temporal development of the traction forces would match the dynamical changes found for the internal monolayer stress. While this was overall the case, we were surprised to observe that internal monolayer stresses exhibited a short lived stress peak immediately after the addition of the chemokine, that relaxed already significantly (*p* = 0.027) at the next measured point (Fig. 3B). This is surprising, as we could not find a similar short lived peak in the average traction forces which kept rising continuously. The amplitude of the first steep rise in internal monolayer stress made up already ca. 75% of the overall increase measured at 20min post treatment (24.0% vs 31.8% relative increase), while for the average traction stress at the first time interval only ca. 25% of the peak increase at 20 min was reached (5.3% vs 21.4% relative increase). Such a short lived, but pronounced peak might be explained by a rapid change in the stress distribution within the monolayer, where after the first increase, some cell-cell contacts are ruptured. The resulting redistribution of the stress transferred over a varying number of cells within the monolayer would then optimize without affecting the absolute traction stress magnitude applied on the substrate at the edges of the pattern. On the other hand this also means that during the first time interval a substantial part of the internal stress increase inside the monolayer can be attributed to longer range transcellular contractions.

### Transcellular stress redistribution is avoided by hydrodynamic prestress

Motivated by these findings, we wondered if a mechanical prestress of the system could effectively avoid the initial peak of the internal monolayer stress. This hypothesis is driven by the idea that the stress transmission between the cells arranges into a homeostasis with minimal variability, to obtain stable stress balance. If a prestress is applied by a fluid flow, as actually is the case for HUVECs in their natural environment, the system should already have adapted to withstand a higher force and hence be less prone to a further stress increase, even when initiated by TNF-*α*. Furthermore, previous studies focusing on fluid shear stress reported an alignment of ECs in the direction of flow accompanied by a remodeling of ECs’ cytoskeleton and their intercellular junctions via mechanotransductional pathways [29–34]. Intercellular stresses have been found to align quickly (from 1 h onward) with the direction of flow and to subsequently attenuate slowly over time [26]. To test our hypothesis, HUVECs were seeded on PAA gels residing at the bottom of a flow chamber, allowed to spread and grow and were finally exposed for about 16 hours to high rates of laminar flow inducing about 1.5Pa (15 dyn/cm^2^) shear stress onto the cells before the actual experiment was conducted. The flow rate was kept this high throughout the measurement (apart from short interruptions during mounting and addition of TNF-*α*). Qualitatively, cells displayed the same response upon TNF-*α* treatment as under no flow and hence zero shear stress conditions regarding the average traction stress (Fig. 3E). However, as hypothesized we find that in this prestressed situation the initial rapid peak that was typically observed for the internal monolayer stress was absent (Fig. 3F). This suggests that prestressing the monolayer will affect the stress transmission within the monolayer. Interestingly, while the amplitude of the peak in average traction stress had about the same level in both conditions, for the average internal normal stress significantly higher peak values could be observed in the presence of flow (ca. 56 % vs. 32% relative increase at 20 min peak, *p* = 0.0022).

### Delayed increase of central traction forces is reduced by hydrodynamic prestress

To get a better understanding of the development of the spatial distribution of both, the traction forces and the internal monolayer stress, we generated kymographs of the radial stress profiles (Fig. 3C,D,G,H). In these kymographs, each horizontal line represents the radial stress profile at the given time, where the color encodes for the normalized stress intensity. At time 0 TNF-*α* was added to the medium. As this application briefly disturbed the measurement, the corresponding images cannot be analyzed (black line). As expected, the radial kymographs show an increase in both traction and internal monolayer stresses after chemokine addition, which is consistent with the overall time evolution of the stresses. The kymographs show that the described rapid increase in traction stress is largely generated at the edge of the monolayer, for both the situation with and without flow induced shear stress (Fig. 3C,G). Surprisingly, a closer inspection of the traction force kymographs for both conditions shows differences in the inner region of the monolayer patch. While in the situation with applied flow and hence shear stress, only a marginal traction stress increase in the monolayer region is found, this is much more pronounced in the situation without shear stress. This difference becomes even more striking when comparing the relative changes of traction forces of edge and inner region in the two conditions (see S2 Fig.). This evident difference in the traction force dynamics in the inner region of the monolayer is especially interesting in the context of the observed rapid internal monolayer stress peak that was basically undetectable in the prestressed situation. Turning to the kymographs of the internal monolayer normal stress (Fig. 3D,H), the rapid increase is detected throughout the inner region of the monolayer and, consistent with the average values, this is absent in the prestressed situation. It is striking to see such a quick modulation of the internal monolayer stress exclusively in the non-prestressed situation, although this behaviour is not mirrored in the traction forces applied at the edge. We suspect that indeed the stress redistribution between the cells is dynamically adapting to the overall load, and that in situations with preload no redistribution is required despite the increase in overall stress on the cells, potentially due to more maturated cell-cell adhesion in the face of shear stress [33, 35, 36].

### Qualitative response of a HUVEC monolayer is independent of TNF-*α* concentration

As this hypothesis suggests that the speed of contraction increase is relevant for such stress redistribution, we wondered if changing the TNF-*α* concentration would allow to modulate the increase of traction forces and internal monolayer stress. To test this we applied three concentrations, namely 5, 20 and 100 ng/ml TNF-*α* to the HUVEC monolayers. As presented in Fig. 4A,B, we could only find an overall delay of the response. However, the actual speed of contraction increase was not affected by the different concentrations. Interestingly, when looking at the rapid intermediate peak in internal monolayer stress, the delay in response seems to make this peak more pronounced in the low TNF-*α* concentration, although the absolute value of the peak is less.

**Fig 4.**
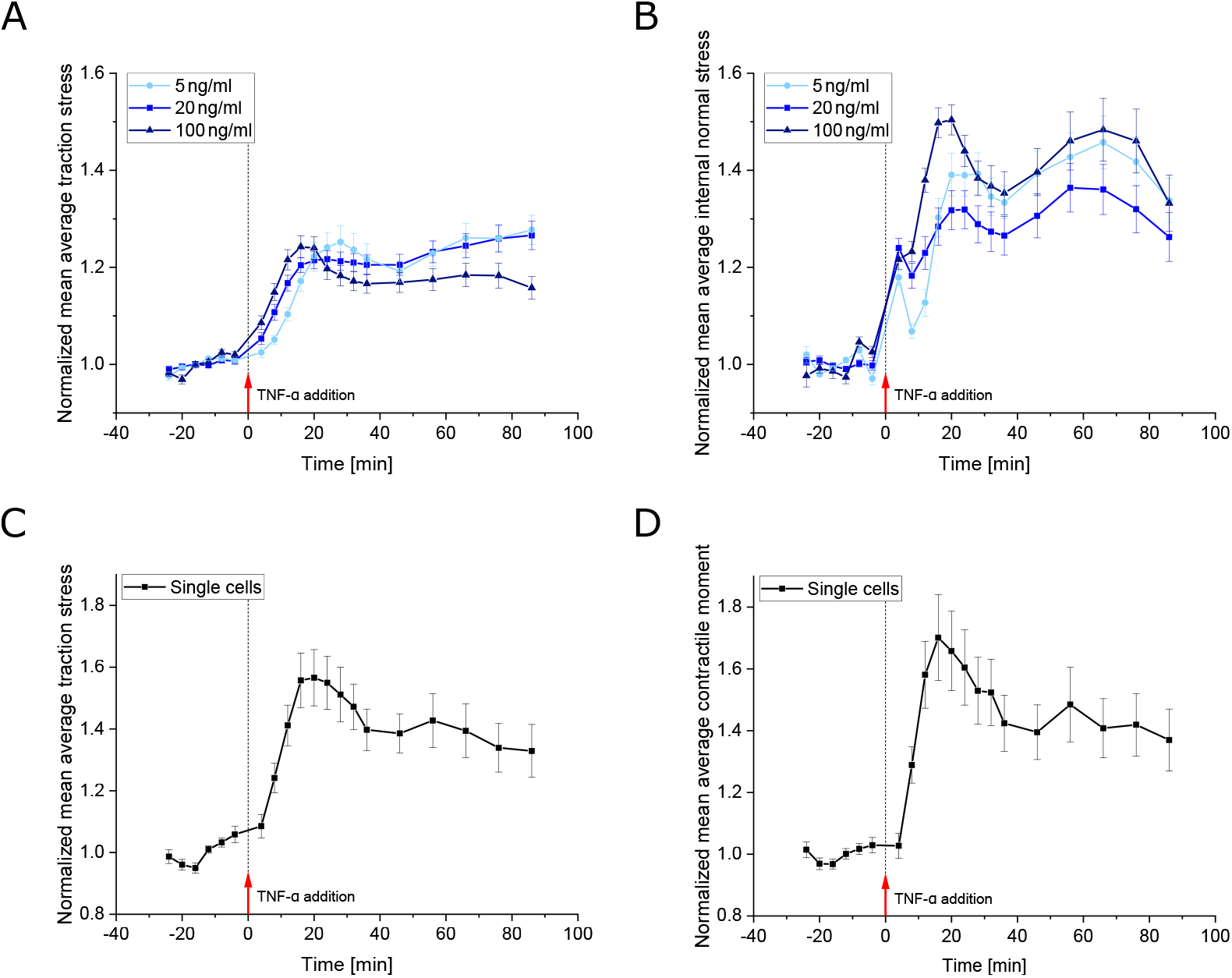
Time evolution of the mechanical response upon TNF-*α* exposure for different chemokine concentrations and for single cells. **A,B**: Time course of the mean of spatially averaged traction stress (A) and internal normal stress (B) of multiple monolayer islands recorded at baseline and upon TNF-*α* treatment for 3 different drug concentrations as indicated. Data is shown as mean±SEM and has been normalized to the mean of the first 6 timepoints recorded prior to drug administration, respectively. For each concentration 3 independent experiments were performed with a respective number of *n*_5ng_ = 29, *n*_20ng_ = 29 or *n*_100ng_ = 33 of monolayer patches. No clear difference in strength of the response can be observed for the different concentrations. However, the response appears to occur slightly faster with increasing concentration, while the drop in internal normal stress after the first steep rise appears progressively reduced. **C**: Same as A, but for single endothelial cells instead of EC monolayers and at 20 ng/ml TNF-*α* concentration only (*n* = 26, *N* = 3). Single ECs display qualitatively the same response to TNF-*α* as EC monolayers while reaching significantly higher peak values (*p* < 0.001 at 20min peak). **D**: Same as C, but showing the contractile moment instead of the traction stress. The contractile moment serves here as a measure of intracellular stress as compared to internal normal stress in the case of monolayers of ECs. However, no initial intermediate peak is found here but the time course rather matches the one observed for the average traction stress.

### Single cell response is attenuated by contact in monolayers

Considering the qualitatively different initial behaviour of the internal monolayer stress, we wondered to which extend collective effects can be separated from the single cell response to TNF-*α*. To this end, we seeded single cells on the same elastic substrates and measured their relative increase in traction forces upon TNF-*α* stimulation at 20ng/ml concentration. As shown in Fig. 4C, we find a qualitatively similar response of single ECs upon chemokine stimulation. However, interestingly, the relative increase of stress was with > 50% significantly (*p* < 0.001) larger than the increase of traction stress measured for the monolayers. However, this difference marginalized after ongoing presence of TNF-*α*. These results suggest that in the monolayer situation, TNF-*α* stimulates an increase of force transmission to the neighbouring cells, which is eventually not visible on the overall average traction force level. A possible explanation for the observed difference could be found in the intracellular signaling in response to cell-cell force transmission, which can only occur within the monolayer. However, the details of such signaling effects need to be further studied. When considering the contractile moment of single cells as a measure of intracellular stress, we find no initial internal stress redistribution to occur within single cells as compared to the observed stress redistribution inside a monolayer on a transcellular scale.

### Contraction, and not actin restructuring is required for transcellular stress redistribution

As the presence or absence of the initial peak in the internal monolayer stress dynamics was found to be dependent on a prestressed situation, we suspected that it occurs indeed due to a reorganization of stress rather than a reorganization of the force transmitting structures such as the actin cytoskeleton. To get further insights into this question, we decided to perturb the intracellular signaling mediated by the small RhoGTPases RhoA and Rac1. While RhoA is generally known to control the activity of myosin motors, and therefore directly affect cellular contractility, Rac1 is known to be a key player in controlling the restructuring of the actin cytoskeleton. On top of that, RhoA and Rac1 have previously been reported to be involved in the formation of stress fibres and other organizational changes in the actin cytoskeleton in HUVECS in response to TNF-*α* already within 15-30 min of stimulation [17]. Further RhoA activity levels have been reported to have yet increased after 1-5 minutes of TNF-*α* exposure [18, 19]. Phosphorylation of RhoA is known to lead to the activation of ROCK which in turn triggers myosin mediated contractility via phosphorylation of the myosin light chain. Hence, it is possible to test this signaling pathway by inhibition of ROCK activity. Consistent with this, the inhibition of ROCK via increasing concentrations of the drug Y27632 led to a reduction of average traction stress settling after about 20 min for progressively lower plateau levels (Fig. 5A). Also the increase of average traction stress after TNF-*α* exposure found for control cells could be observed successively less with higher concentrations of ROCK inhibitor and ended up essentially undetectable. Without any obvious differences, ROCK inhibition had the same effect on the internal normal stress (Fig. 5B), with the exception of the initial steep rise in internal monolayer stress in response to TNF-*α*, which was fully absent. This was also the case for 10 *μ*M inhibitor concentration, where we could still observe an overall relative increase in traction stress as well as internal normal stress after 20 min of TNF-alpha stimulation (*p* = 0.018 and *p* = 0.033, respectively, for 20min compared to −4 min).

**Fig 5.**
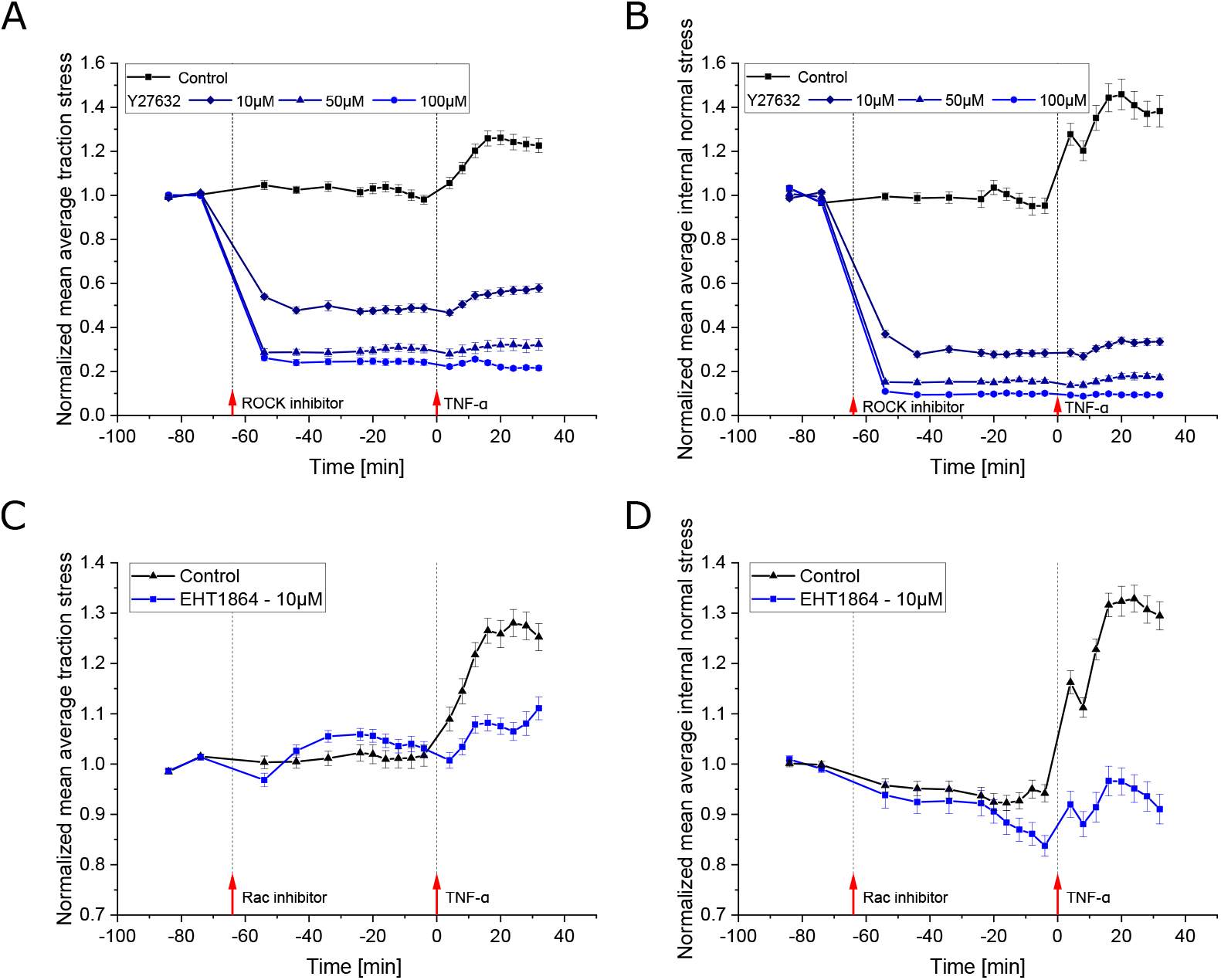
Effect of ROCK and Rac inhibitors on the mechanical response of HUVEC monolayers upon TNF-*α* stimulation. **A,B**: Time evolution of the mean of spatially averaged traction stress (A) and internal normal stress (B) of multiple monolayer islands for control condition and 3 different concentrations of ROCK inhibitor (Y27632). Stresses were recorded at baseline, for about 1hour of ROCK inhibitor treatment and upon 20ng/ml TNF-*α* stimulation. Data is shown as mean±SEM and has been normalized to the mean of the first 2 timepoints recorded prior to drug administration, respectively (*n*_ctrl_ = 12, *n*_10*μ*M_ = 10, *n*_50*μ*M_ = 7, *n*_100*μ*M_ = 11). Inhibitor treatment causes average stresses to drop by more than 50 %, reaching successively lower plateau levels with increasing drug concentrations. The response to TNF-*α* is progressively attenuated and rendered essentially undetectable for the highest concentration. The initial peak in internal normal stress observed in control condition is lost already for the lowest inhibitor concentration applied. **C,D**: Same as A and B, respectively, but for the application of Rac1 inhibitor (EHT1864) instead of ROCK inhibitor and for 10 *μ*M concentration only. Three independent experiments were conducted for both control and drug condition, both recorded on the same day for the same batch of cells, respectively (*n*_ctrl_ = 33, *n*_10*μ*M_ = 33). Average stress levels are only slightly disturbed by Rac1 inhibitor treatment. The response to TNF-*α*, however, is reduced to a similar amplitude as in the case of ROCK inhibitor pre-treatment.

We next turned to the inhibition of Rac1 via EHT1864. Rac1 is known to modulate the actin cytoskeleton via the WAVE/Arp2/3 pathway [37–39]. The inhibition of Rac1 led to only a small disturbance in the average traction stress level ending up slightly higher as compared to control conditions before stimulation with TNF-*α* (Fig. 5C).

Interestingly, when adding TNF-*α* after pretreatment with the Rac1 inhibitor, the rise in average traction stress observed in control situation was suppressed to a similar degree as was the case for ROCK inhibition at the same concentration. When considering the lower level of traction stress in case of ROCK inhibition prior to TNF-*α* treatment though, the relative increase following TNF-*α* stimulation was even more attenuated in case of Rac1 inhibition. Further, the response appeared slightly delayed as compared to control situation. The time evolution of the internal stress response to TNF-*α* on the other hand closely matched with control condition. The amplitude of the response though was reduced by a similar amount as observed for the average traction stress (Fig. 3D). Even when perturbing the signaling responsible for the restructuring of the actin cytoskeleton, we could still observe the initial peak in internal monolayer stress, suggesting that indeed the redistribution of stress, and not a redistribution of the actin cytoskeleton is responsible for the fast dynamics of the internal monolayer stress.

## Results complement former biological observations

Former studies of early TNF-*α* stimulation by Wojciak-Stothard et al. [17] reported prominent stress fiber formation in HUVEC monolayers after 15-30 minutes of TNF-*α* exposure. This timing coincides with our observation of a peak in increased traction and internal monolayer stresses in response to TNF-*α* after around 20 min. Stress fiber formation has been found to follow increased MLC phosphorylation initiated by elevated levels of RhoA activity [18, 19, 40]. While RhoA activity was reported to be raised at 1-10 minutes of TNF-*α* exposure, enhanced levels of phosphorylated MLC were to our knowledge not measured before 30 minutes, so it remained questionable how quickly actomyosin contractility increases upon TNF-*α* treatment. Our results show a rapid increase in EC monolayer stress occurring immediately (within our time resolution) after stimulation suggesting quick downstream processes following RhoA activity. While substrate traction levels increased continuously, the internal normal stress rose most intensely within the first measured time frame after exposure followed by a small intermediate drop, most likely explained by alterations in transcellular force transmission. Further, the internal normal stress levels reached a slightly higher overall increase than was the case for substrate traction. All together reflect a particular increase of cell-cell forces. The latter fits indications from the work of Millan et al. [41] who found stress fiber ends to connect discontinuous adherens junctions from neighboring cells in response to TNF-*α*. Also they observed no increase in the number of focal adhesions - the predominant structures to transmit contractile forces of stress fibers to the substrate - although the number of stress fibers increased upon TNF-*α* stimulation. It should be noted though, that their observations were made 20 hours post treatment. Assessing TNF-*α* signaling through inhibitor application, we found that ROCK inhibitor Y27632 treated cells showed successively less up to hardly any reaction to TNF-*α* in neither substrate nor cell-cell forces with increasing inhibitor concentrations. As RhoA is known to act downstream via ROCK, this finding underpins the role of RhoA in the mechanical response to early TNF-*α* stimulation. Besides RhoA also Rac1 was acknowledged to be essential for early stress fiber formation in response to TNF-*α* [17]. In general, however, RhoA and Rac1 are known to have opposite effects on endothelial barrier function [42–44] and have even been found to be linked in a double-negative feedback loop in mesenchymal breast cancer cells [45]. Applying the Rac1 inhibitor EHT1864 onto our EC monolayer islands attenuated the response to TNF-*α* treatment to a similar level as was the case for ROCK inhibitor treated cells at the same concentration. Taking into account the prior significant drop in overall stress level observed in case of ROCK inhibitor treated cells as opposed to the only small disturbances caused by the Rac1 inhibitor, the relative increase in traction and internal monolayer stress upon TNF-*α* stimulation was even more dampened following Rac1 inhibition. Despite the known antagonistic effects of RhoA and Rac1, these findings underline the importance of Rac1 for the enhancement of contractile machinery in ECs following TNF-*α* stimulation.

## Conclusion

In the last decades endothelial cell mechanics have been established to play an important role in many inflammatory processes. In this study we uncovered the mechanical stress response of HUVECs in the particular case of early TNF-*α* stimulation. In our in vitro assay 300 *μ*m diameter monolayer EC islands were cultured on PAA gels of 15 kPa stiffness. High traction stresses appeared at the edges of the monolayer patches and were found to be transferred to a substantial degree to neighboring cells increasing monolayer tension towards the patches’ center. This finding is in agreement with observations reported for other monolayer systems like epithelial cells [27]. Administering TNF-*α* to the cells, we found a rapid increase in average substrate traction as well as internal monolayer stress rising quickly towards a typical peak after about 20 minutes. The peak timing coincides with former observations of prominent stress fiber formation [17]. While substrate traction levels increased continuously, the internal normal stress rose most intensely within the first measured time frame after exposure followed by an intermediate drop. This suggests longer range transcellular contractions and a rapid change in stress distribution within the monolayer. We hypothesize that after the first increase some cell-cell adhesion might rupture and that stresses redistribute subsequently without affecting the absolute traction stress magnitude applied on the substrate at the edges of the pattern. Mechanically pre-stressing the cells via flow induced shear stresses, we found the described intermediate peak in internal normal stress to be abrogated. Further, the overall increase in internal normal stress was higher than in condition without flow. We speculate that in situations of external pre-stress no stress redistribution is required and cells can withstand a higher overall mechanical stress increase, potentially due to more maturated cell-cell adhesion in the face of shear stress. Thus, investigations further unravelling the intercellular adhesion dynamics would be of great interest.

While TNF-*α* is mostly known for its longer timescale role in enhanced permeability and sustained leukocyte recruitment after several hours post stimulation, the rapid mechanical response reported here might hint to a possible role of enhanced contractile machinery to support leukocyte recruitment early on. Another possibility is that the increase in monolayer tension is simply a self stimulating necessity to induce the previously observed actin cytoskeletal changes that would then become relevant in the long run for the aforementioned enhanced permeability and sustained leukocyte recruitment.

## Materials and methods

### Cell culture and treatments

Human umbilical vein endothelial cells (HUVECs) were a kind gift from Prof. Dr. V. Gerke (Institute of Medical Biochemistry, University of Münster, Germany). The cells were cultured in 60 mm diameter polystyrene dishes (Corning CellBIND Surface 60 mm Culture Dish, Corning) in culture medium consisting of two equal parts of Endothelial Cell Growth Medium 2 (PromoCell) containing its according supplements (Growth Medium 2 SupplementMix, PromoCell) and of Medium 199 Earle’s (F0615, Biochrom GmbH or M2154, Sigma-Aldrich) with 2.2 g/l NaHCO_3_, without L-glutamine supplemented with 10% fetal bovine serum (F7524, Sigma-Aldrich), 0.2 units/ml heparin (H3149, Sigma), 30 μg/ml gentamicin (gibco) and 15 μg/ml amphotericin B (gibco) as previously described [46]. Culture dishes were placed in a humidified incubator at 37° and 5%CO_2_. Cells were used for experiments at passages 3 to 6.

For staining, cells were treated with CellMask Deep Red plasma membrane stain (Thermo Fisher Scientific). Drugs were applied at indicated concentrations and time. ROCK 1&2 inhibition was performed via Y-27632 -dihydrochlorid (Sigma-Aldrich) and Rac1 inhibition using EHT1864 (MedChemExpress). Recombinant human tumor necrosis factor-*α* was purchased from gibco.

### PAA gel preparation with patterned coating

Polyacrylamide gels of 15 kPa stiffness (confirmed by rheological measurements and nanoindentation) were prepared as follows. To ensure stable attachment of gels to the surface of either a glass bottom dish (CELLVIEW - 35mm, Greiner Bio-one International) or a glass coverslip for sticky-Slides (ibidi), the latter were first cleaned with 0.1 N NaOH, treated with (3-Aminopropyl) trimethoxysilane (APTMS) for 3 minutes, thoroughly washed and then incubated with 0.5 % glutaraldehyde solution for 30 minutes.

To create patterned surface coating of gels, 12 mm diameter round glass coverslips were first cleaned with a plasma cleaner (UV Ozone Cleaner, BioForce Nanosciences) and then incubated with 0.1 mg/ml poly(L-lysine) poly(ethylene glycol) co-polymers (PLL-PEG, SuSoS) in 10 mM HEPES solution for 1 hour and washed afterwards with double-distilled water. With the help of an anti-reflective chrome coated Quartz-glass photomask (Delta Mask B.V.) circular holes of 300 μm diameter were burned into the PLL-PEG layer using UV radiation. After quick washing treated sides of the coverslips were then covered for 1 hour with 90 μg/ml collagen type I (Corning Collagen I, Rat Tail, Corning Life Sciences).

A gel premix was prepared by first mixing 500 μl of 40 % acrylamide solution with 250 μl of 2% N,N’-Methylenebisacrylamide solution. Out of that solution 113 μl were mixed with 372 μl of 0.65X PBS and 15 μl of fluorescent bead solution (100 nm diameter, NH_2_ coated micromer-redF, Micromod). Polymerization of gel premixes was started via 10% ammonium persulfate solution (APS) and 1.5 μl of N,N,N’,N’-Tetramethylethylenediamine (TEMED). Following quickly, 5 μl of polymerizing premix solution were put onto the surface of the glass bottom dish and covered with the pre-patterned round coverslips.

For flow experiments polymerizing gel solution was placed on a glass coverslip with dimensions to later fit on a bottomless channel slide (sticky-Slide I Luer, ibidi). Two tesafilm stripes were temporarily attached to the glass coverslip beforehand and the pre-patterned top coverslip put on top of them to create a channel with defined height and width to restrict the drop of gel solution in a way to later properly fit into the channel of the resulting flow chamber after sticking the glass coverslip to the bottom of the sticky-Slide.

All chemicals used were purchased from Sigma-Aldrich (Merck) if not stated otherwise.

### Experimental procedure and microscopy

For all experiments cells were seeded out 18-24 hours before measurement. For experiments without flow and confluent monolayers of cells, 200 μl of 1.6 × 10^6^ cells/ml containing solution were seeded out into 1 ml EC medium in a 35 mm diameter glass bottom dish. For single cell experiments 3 × 10^4^ cells were seeded into the same kind of dish, respectively. After 1 hour samples were washed once to remove dead cells and refilled with 2-3 ml medium. For flow experiments the channel of the flow chamber slides was filled with 150 μl of 1.6 × 10^6^ cells/ml containing EC medium. Low levels of flow were applied after 2-3hours resulting in about 0.15Pa (1.5dyn/cm^2^) and 0.5Pa (5dyn/cm^2^) shear stress for 1 hour each. After that cells were exposed to high shear stress of about 1.5 Pa (15dyn/cm^2^) until the start of the measurement. Samples were kept in a cell culture incubator at 37° and 5%CO_2_ until measurement.

For microscopy, samples were mounted inside a small stage top incubator (UNO Top Stage Incubator, H301-Mini, OKOLAB) on an inverted microscope (Nikon Eclipse Ti-E) and kept there again at 37^°^ and 5 % CO_2_. The microscope was equipped with a spinning disk head (CSU-W1 Yokogawa). Images were acquired via a scientific CMOS camera (Prime BSI, Photometrics) using the Slidebook 6 software (Intelligent Imaging Innovations). Using a 40X water-immersion objective (CFI Apo LWD Lambda S 40XC WI, Nikon) with a numerical aperture of 1.15 time-lapse measurements were performed recording 3D stacks of fluorescent beads inside the gels with an excitation laser of 561 nm wavelength and of ECs using bright field illumination. For experiments on plasma membrane stained cells a 60X water-immersion objective (CFI Plan Apo VC 60X WI, Nikon) with a numerical aperture of 1.2 and an excitation laser of 647 nm wavelength for EC images was used instead. In this case, to record the full monolayers, four 3D image stacks at four different positions with about 25 % overlap were recorded for each patch and stitched together using a custom ImageJ-macro (based on [47]). Z-planes were recorded in all cases with 6 μm range a 0.33 μm interval centered around the plane most close to the gel’s surface that contained fluorescent beads. Drugs were applied by aspirating 2/3 of the medium from the sample dishes mixing it with the drug and readding it. In the case of flow experiments drugs were added to the syringe reservoirs after pausing the flow, mixed with the medium with a big pipette and applied by restarting the flow.

At the end of each time-lapse acquisition 0.5 ml of 5 % Sodium dodecyl sulfate (SDS) solution were added to disintegrate the cells and another timepoint was recorded as reference for no deformation.

### Traction force and monolayer stress derivation

Calculations were performed similar as previously reported [48]. Traction forces were derived from PAA gel deformations based on displacements of fluorescent beads close to the gel’s surface with respect to a reference state without any cell induced deformations. The reference state was created by lysing the cells via a SDS solution. Displacements were determined by an iterative free form deformation algorithm carried out with the open source software Elastix [49]. In short, the 3D reference image was iteratively deformed via B-spline functions with central nodes starting as a regular mesh and compared to the 3D image of a respective timepoint that contained potential gel deformations. Optimizing the quality of agreement between both images measured by the advanced Mattes mutual information metric the deformation was tuned in each iteration accordingly following an adaptive stochastic gradient descent algorithm. The calculation went down in a three level pyramid approach from coarse to finest scale dividing the grid size in half and doubling the number of iterations for each level with 1.3 × 1.3 *μ*m grid size and 2000 iterations at the finest scale. The number of random samples per iteration was chosen with respect to the size of each image. Finally, plugging the resulting deformation field into the equation for the Tikhonov regularized elasticity problem for finite thickness substrates and solving it in the Fourier domain [50, 51], 3D traction forces exerted onto the gel’s surface were derived using a custom-made MATLAB (MathWorks) program. Picking up the idea of Huang et al. [52], Tikhonov regularization was performed combining Bayesian theory with an estimation of background variance to provide a less subjective and more stable choice of the regularization parameter.

For further evaluations the area covered by the cells was determined. In the case of stained EC monolayers, the region was inferred from the fluorescence signal via thresholding. For unstained EC monolayers active contours were applied to a manually created outline based on the 2D in-plane traction force field. The validity of the resulting monolayer outline was checked with the help of the recorded bright field images of the cells. As for single cells the aforementioned approach led to significant discrepancies due to the discontinuity of the force distribution at the cells’ edges, the covered region was manually created based on the bright field images and slightly locally extended to include force spots that were not fully lying within the cell’s area due to imperfections of the force derivation. To do so, bright field images were reversely deformed according to the determined deformation fields to match the reference frame of the calculated traction fields and overlaid with an image of the latter.

Evaluation of average traction stress values was restricted to the cell or monolayer region.

This region - created as described - was further used in the case of monolayers for the derivation of the internal stress field. Extruding this region into 3D with 5 μm height an object was created for finite element analysis modelling the EC monolayer as a homogenous, linear elastic material. Imposing the beforehand derived traction forces (only considering 2D in-plane forces) as boundary load to the bottom plane of this object, the internal stress tensor was inferred via force balance using finite element calculus performed with the COMSOL Multiphysics software package as previously described [48]. Settings used: boundary condition ‘rigid motion suppression’, material property ‘nearly incompressible’, automatically created tetrahedral mesh with ‘finer’ size. To deal with small rotational artifacts arising locally around a position that depended on the initial orientation of the object, the analysis was performed for four different orientations of the object in-plane rotated by 90° each. For each rotation the region exhibiting clear artifacts was omitted and the final result obtained as an average of the results for all rotations. The results presented in this work solely focus on normal stresses rather than shear stresses and are given as an average of the xx- and yy-component of the derived stress tensor field.

### Flow setup

For flow application the ibidi pump system was used. It consists of an electronically fine tuned air pressure pump and a fluidic unit that holds two syringes and houses a valve that regulates the flow through the tubing. The air is taken in from a 5 % CO_2_ incubator and pumped to the syringes which are connected over intertwined tubing to a flow chamber slide in a way that yields unidirectional flow through the latter while transferring fluid from one syringe to the other back and forth switching between valve states. As mentioned above, bottomless chamber slides (sticky-Slide I Luer, ibidi) were stuck on a glass cover slip to create a flow chamber slide with 0.65 mm height, 5 mm width and 50 mm length. The PAA gels residing on the glass coverslip within the chamber had varying heights of 54-72 μm. The shear stress τ acting on cells at the gels’ surface was estimated via the following equation [53]:

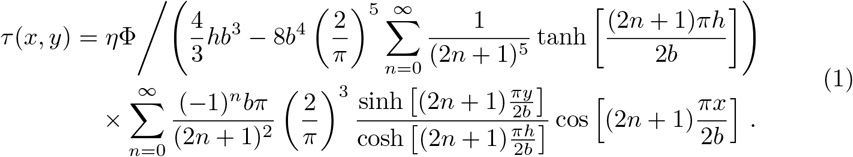

Here *η* is the viscosity of the medium, Φ the total flow through the channel, 2*h* the height and 2*b* the width of the channel. The coordinates *x* and *y* give the position centered with respect to the channel, meaning *x, y* = 0 at the center, *y* = – *h* at the bottom of the channel. The gel inside the chamber reduces the effective channel height to 578-596 μm. Although our gels are only 4.5 mm instead of 5 mm in width, the difference in local flow due to this variance in cross section area is less than 1.2 %. Assuming a viscosity of *η* = 0.72 mPa o s for 37°C warm medium, the shear stress experienced by the cells at the surface of the gel lies for the highest applied flow rate of Φ = 35.66ml/min between 1.46 and 1.64Pa (14.6-16.4dyn/cm^2^) dependent on the actual gel height and the position in *x*.

### Statistical analyses

Statistical comparison between groups of data was performed either via a two-tailed *t*-test or a Mann–Whitney–Wilcoxon test depending on if the data would qualify as normally distributed at 5 % confidence level according to a Shapiro-Wilk test. In case either or both of the two compared data sets failed to be significantly drawn from a normal distribution, the Mann-Whitney-Wilcoxon test was employed to evaluate the significance of potential differences between the compared data sets. The latter was the case for all comparisons performed with the exception of the data shown in Fig. 2B, the initial rise at 4 min in traction forces (Fig. 3A), the increase at 20 min with respect to −4 min in case of ROCK inhibitor treated cells after addition of TNF-*α* and the data shown in S2 Fig. Number of analyzed individual monolayer patches and number of independent repeats performed is stated in each figure caption, respectively.

## Acknowledgments

We gratefully acknowledge the support of the Cells in Motion Cluster of Excellence (EXC 1003 CiM) and the financial support of the European Research Council (ERC-CoG, PolarizeMe 771201). We also thank the Interdisziplinäres Zentrum für Klinische Forschung (IZKF, Bet1/013/17) of Münster.

