## Supporting information for "Human endothelial cells display a rapid and fluid flow dependent tensional stress increase in response to tumor necrosis factor-*α*"

**S1 Video.**
**Membrane stained cells display contraction upon TNF-$\alpha$ exposure.** Image sequence of a representative plasma membrane stained HUVEC monolayer island about 1 h before and 1 h after the addition of TNF-$\alpha$.

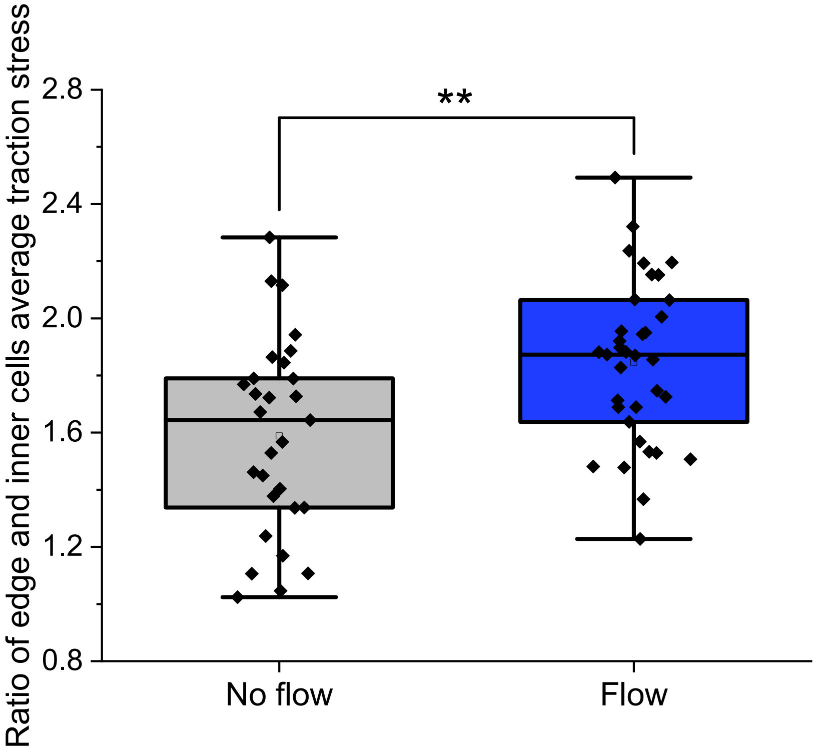


**S2 Fig. Ratio of edge and inner cells' average traction stress for flow vs. no flow condition at peak time.** Comparison of edge by inner cells average traction stress ratios for flow (1.5 Pa shear stress) and no flow (zero shear stress) experimental conditions evaluated at peak increase at 20 min past TNF-$\alpha$ addition reveals significant difference. $n_{\text{no\_flow}} = 29, n_{\text{flow}} = 35, N_{\text{both}} = 3$. **, $p < 0.01$ ($p = 0.0016$).
