## Supplementary figures and images for "Human endothelial cells display a rapid and fluid flow dependent tensional stress increase in response to tumor necrosis factor-*α*"

### S1 Video

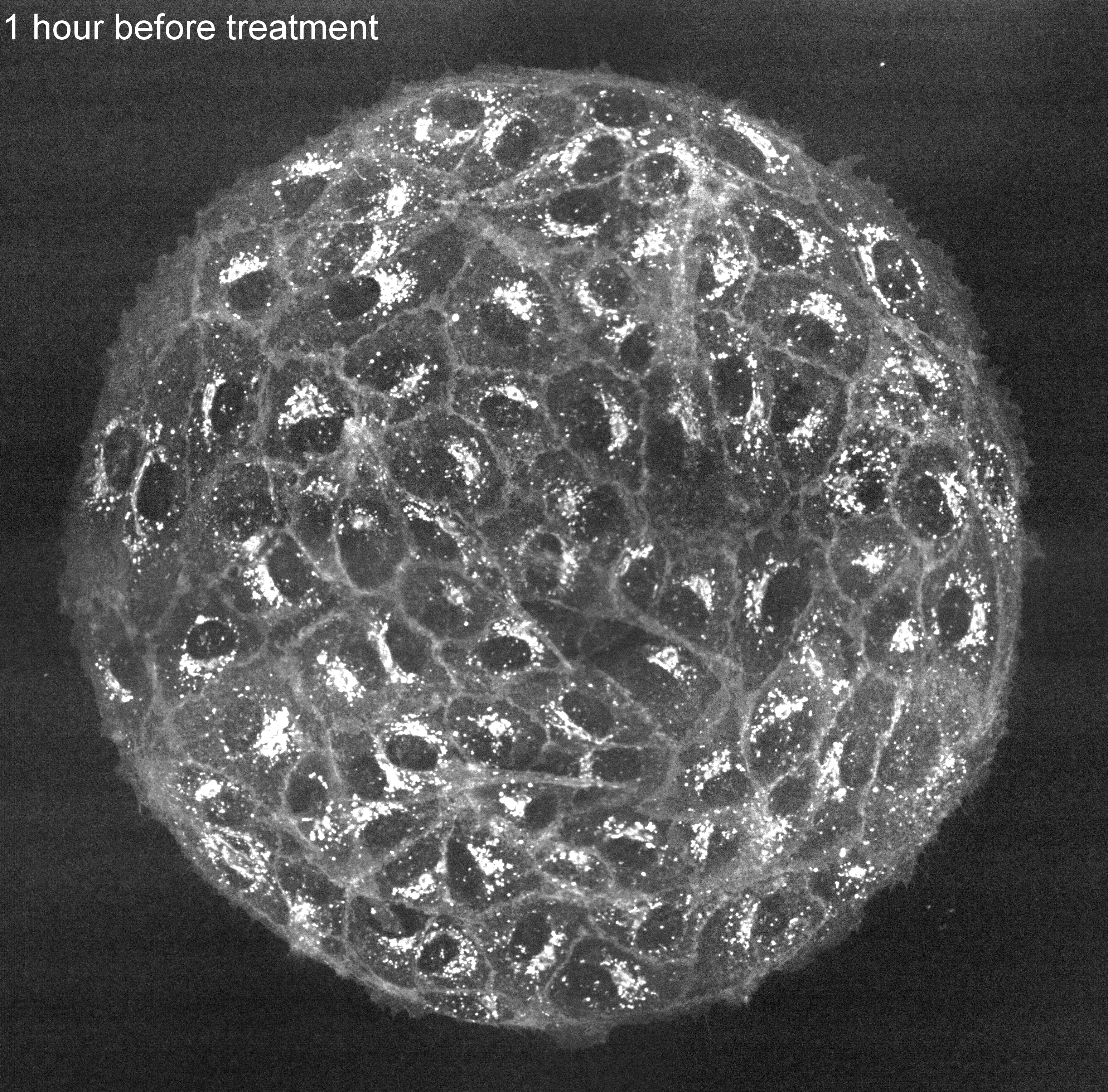
